# Hemifield-Specific Motion Extrapolation Reveals Limits of Interhemispheric Integration

**DOI:** 10.1101/2025.10.06.680800

**Authors:** Coleman E. Olenick, Mazyar Fallah

## Abstract

The accurate perception of moving objects is a fundamental challenge for the visual system, which must compensate for neural processing delays. Motion extrapolation is a proposed mechanism whereby the brain predicts an object’s future position. We investigated how an object’s motion history shapes its perceived position using the flash-jump illusion, in which a brief color change in a moving bar is mislocalized as further along the direction of motion. Across two experiments, we found that longer preceding motion sequences improved localization accuracy, consistent with motion adaptation. This effect occurred regardless of whether motion continued after the flash. Notably, mislocalization transiently *reappeared* as the object crossed the vertical midline, suggesting that motion adaptation and motion extrapolation operate independently between within each hemifield. Manipulating the length of the sequence in each hemifield in Experiment 2 confirmed that this adaptation is spatially confined to each hemifield, with limited interhemispheric transfer. The results align with a Bayesian framework in which the brain integrates signals from both hemispheres, with midline crossings triggering a shift from adapted to unadapted neural populations. We identify motion extrapolation, supported by hemifield-specific adaptation in area MT and integration in area MST, as the mechanism behind these midline discontinuities. This work reframes smooth pursuit not just as a tracking behaviour, but as a functional solution to overcome the inherent limits of interhemispheric motion processing.

## Introduction

Accurately perceiving and interacting with moving objects is a fundamental challenge for the visual system. Given the inherent delays in neural processing that arise from the time required for sensory information to travel through the visual hierarchy, misalignments arise between the brain’s representations and the real positions of objects in the world. To compensate for these delays, the visual system must employ mechanisms to ensure accurate localization of moving objects. For instance, when driving at high speeds, drivers must anticipate the movements of other vehicles and pedestrians. If neural processing were purely reactive, drivers would consistently misjudge positions, yet they are often able to respond appropriately by predicting where objects will be in the near future. Motion-related perceptual illusions have been employed extensively to study the ways the brain compensates for objects in motion (Hogendoorn, 2020). Emerging from this research is the theory of motion extrapolation, which posits that the brain makes predictions about the motion of an object that are used to shift its processing further along its trajectory through the visual hierarchy (Turner et al., 2024).

A particularly influential paradigm for the study of motion extrapolation is the flash-lag effect (Nijhawan, 1994). In the flash-lag experiment, a bar is moved across a screen at a constant speed and a stationary bar briefly flashed adjacent to the moving bar is perceived as having flashed behind the moving bar. This finding has led to the development of various manipulations in flash-related motion illusions (Hogendoorn, 2020). One such manipulation, the flash-jump illusion, removes the physical separation between the stationary flash and the moving bar, instead briefly changing the color of the moving bar. Observers of the flash-jump illusion reporting the flash later than it actually occurred, a result dubbed the flash-jump effect (FJE) (Sundberg et al., 2006; Eagleman and Sejnowski, 2007). Recent work on the FJE has demonstrated that the color of the flash can modulate the degree of extrapolation in a manner consistent with an attentional weighting of the actual flash position in a Bayesian perceptive framework (Saini et al., 2021).

Applying Bayesian principles to motion-induced position shifts is well-established (Koechlin et al., 1999), with their association becoming more widespread in recent years. Computational models have been developed with a Bayesian basis to explain the flash-lag effect (Khoei et al., 2017) and motion perception more generally (Yang et al., 2021; Jiang and Rao, 2024). The central principle of Bayesian approaches to motion perception is that the brain combines noisy sensory inputs with prior expectations about the motion of an object to form a posterior estimate about that object’s position (Weiss et al., 2002). Internal models of motion extrapolation are formalized as priors that are constantly updated by recent object motion and improved by coherent motion (Rao et al., 2004). When sensory evidence is weak, noisy, or otherwise unreliable, object localization relies more heavily on these motion extrapolation mechanisms (Stocker and Simoncelli, 2006; Khoei et al., 2017). The Bayesian perspective provides a unifying framework for understanding motion-induced position shifts, including the flash-lag effect, FJE.

We are interested in how an object’s motion history affects motion extrapolation. How does the length of time that an object has travelled along a given trajectory influence the localization of that object at a specific moment in time? Early work on the duration of the moving bar was done using the flash-grab illusion, in which a stationary object is flashed at the exact moment a moving background reverses direction, and is perceived to be pulled or “grabbed” in the new direction of motion. This work found conflicting results, reporting an increase (Vreven and Verghesse, 2005) or decrease (Bachmann et al., 2003; Cantor and Schor, 2007) in the size of the flash-lag effect. However, Linares and colleagues (2007) demonstrated that the influence of duration was modulated by the distance between the moving bar and the flash. Specifically, they found that the flash-lag effect increased with the duration of the pre-flash trajectory only when the flash was presented far from the moving object; the duration had little influence when the flash was presented nearby. More recent work has found that the flash-grab effect is increased for longer preceding motion history (Coffey et al., 2019; Takao et al., 2022). Coffey et al. (2019) attributed the increased illusion to an increased expectation of the flash within the experimental windows. These results create a spatial separation between either the flash and the preceding motion, or in the case of Coffey et al. (2019), a spatial discontinuity between preceding and succeeding motion via the flash-grab effect. In this study, we examine the effects of motion history on motion extrapolation without spatial separation of the flash from its motion history by using the FJE. Based on prior results, we expect that longer preceding motion sequences would result in an increase in the effects of motion extrapolation producing an increase in the FJE, but only when there is motion after the flash (Sundberg et al., 2006).

### Experiment 1

#### Methods

Note that the methods and data from experiment 1 have been previously published by Saini et al., (2021), but the data are being used for the novel analyses presented here.

##### Participants

Twenty-four participants volunteered for this study, but were naïve to the purpose (7 Males, range: 18-43 years). Ethics approval was obtained from the York University Human Participants Research Committee. Written consent was acquired prior to participation, and colour vision was ensured using the Ishihara test for color blindness (Ishihara, 2006).

##### Apparatus

Participants were sat in a dimly lit room with their head placed on a chinrest for the duration of the experiment. The chinrest was positioned 84cm away from a 21” CRT monitor (60Hz, 1024×768). Throughout the experiment, participants were required to maintain a fixation, which was monitored using an Eyelink II infrared eye tracker (SR Research, Ontario, Canada) to track the right eye position. The experimental procedure and data collection was programmed using Presentation software (Neurobehavioral Systems, Inc., Berkeley, CA).

##### Procedure

At the start of the experiment the eye position was calibrated, and throughout as required. The trials began with the participant fixating a grey cross (0.23° X 0.23°; CIExy = [10.83, 16.25]; luminance = 11.93cd/m^2^) presented on a black background (CIExy = [0.35, 0.56]; luminance = 0.365cd/m^2^) for 200ms. Following fixation, a light grey bar (0.2° X 2.24°; CIExy = [10.83, 16.25]; luminance = 11.93cd/m^2^) appeared 2.71 dva below fixation and 5.22 dva to either the left or right side of the center of the display. The bar appeared for 2 frames (33 ms), then disappeared for 2 frames, subsequently reappearing 0.77° leftwards or rightwards. This sequence generates apparent motion of the bar with a speed of 11.41 °/s moving left or right from the periphery over 21 bar positions.

During each trial, one of the grey bars instead appeared coloured (luminance = 12cd/m^2^; green: CIExy = [6.007, 2.516]; blue: CIExy = [27.08, 139.3]; red: CIExy = [25.57, 1.804]; yellow: CIExy = [12.24, 2.379]) at 1 of the central 11 bar positions. The motion sequence could either terminate with the appearance of the colored bar or continue after as grey bars again, yielding the terminating and continuing motion path conditions respectively. Once the motion sequence of the bar had completed, participants were asked to indicate the position where the coloured bar had appeared using a probe identical to the grey bars moved with a computer mouse. The probe was locked to the vertical position of the bars but could be moved freely horizontally, with its initial position randomized across the entire range of bar positions. During this response period, participants were permitted to move their eyes freely.

##### Data Analysis

To quantify the magnitude of the flash jump effect (FJE) for a given trial, the horizontal difference between the reported bar position and its actual position was calculated in dva, such that positive values reflect errors further along in the direction of the apparent motion of the bar. The FJE was collapsed across the motion direction of the bar and the 4 colours as those have been previously reported by Saini and colleagues (Saini et al., 2021). The median for every participant was calculated in each of the flash positions and motion sequence termination conditions. The precision of the spatial localization of the illusory flash location was quantified on a participant-by-participant basis by calculating a 95% confidence interval at each experimental condition for each participant, a similar method to those used in reach and gaze paradigms (Blohm and Crawford, 2007; Henriques et al., 2003), and by Saini et al. (2021). A Linear Mixed-Effects (LME) regression model was used to investigate the combined effects of sequence termination conditions and flash position on the FJE magnitude and variability. An LME model allows for a comparison of variation across individual participants, while accounting for general differences in magnitude of the FJE (Meteyard and Davies, 2020). All LME comparisons were made through likelihood ratio tests on maximum-likelihood fit models, with general model significance assessed through comparisons with a null model (only random and fixed intercepts).

### Experiment 2

#### Methods

##### Participants

A new set of twenty-four naïve participants were recruited for experiment 2 (3 Males, 18-27) and were compensated $15 for their time. Ethics approval for the study was obtained from the University of Guelph research ethics board. Written consent was acquired prior to participation, and colour vision was ensured using the Ishihara test for color blindness (Ishihara, 2006).

##### Procedure

Experiment 2 followed a similar procedure with key experimental manipulations to better explore the effects of flash position. Participants were seated in a dimly lit room and positioned with a display 57cm from their eye. The position of the eye was calibrated at the start of the experiment and throughout as required. Like experiment 1, trials began with the participant fixating a grey cross (0.4° X 0.4°; CIExy = [12.28, 18.26]; luminance = 13.49cd/m^2^) on a black background (CIExy = [0.361, 0.5792]; luminance = 0.364cd/m^2^) for 200ms. However, the position of the fixation cross varied between blocks, and was presented at either the center of the display or 4 dva to the right or left of the center for the duration of a block.

After the required fixation, a grey bar (0.34° X 4.14°; CIExy = [12.28, 18.26]; luminance = 13.49cd/m^2^) appeared 7.76 dva below fixation and 11.35 dva to the left or right edge of the display center (measured to the center of the bar), and followed a faster apparent motion sequence than experiment 1 at a speed of 16.82°/s. Along the motion path of the grey bar, one of the bars instead flashed green (CIExy = [6.792, 2.781]; luminance = 13.49cd/m^2^) at 1 of the central 13 bar positions, with the motion sequence continuing or terminating after the green bar. The broader flash and fixation positions allows for a more comprehensive examination of the effects of flash position on FJE magnitude across the visual field from 9.5 dva before to 9.5 dva after fixation in the motion path. Much like the first experiment, motion will occur in both leftward and rightward directions but, given the lack of direction effects in the first experiment, our analysis will collapse the directions. This paradigm also allows for the examination of fixations that occur at early, intermediate, and later times of the overall motion sequence (e.g. an early fixation could be left of center with rightward motion or right of center with leftward motion)

Each participant completed 6 blocks in total repeating each fixation cross position twice. The order of the fixation position between blocks was a randomized sequence of the 3 positions for each participant. Each block contained 52 trials consisting of one trial for each of the 13 flash positions, 2 motion directions (left vs. right), and 2 motion path conditions (terminating vs. continuing).

##### Data Analysis

The FJE magnitude and precision were computed in the same manner as experiment 1 and converted into dva. LME regression models were used to assess the effects of motion sequence length, position of fixation, a binary effect of the flash occurring before or after the midline, and sequence termination conditions on the FJE magnitude and precision. All LME models allowed for participant-based intercepts for each sequence termination condition where relevant, reflecting individual overall responses to the FJE. Follow-up LME models were fit for each fixation position in the flash-continuing sequence conditions, which additionally compared with nonlinear regression was used to compare a linear decrease in FJE magnitude. The nonlinear decrease was represented with an exponential decay for the flashes occurring before the midline: *a*·e^(-*b*·Eccentricity)^ + *c*, where *a* represents the value of FJE at the visual midline, *b* is the decay constant, and *c* reflects the lower limit of the decay and was allowed to vary by participant. The residuals from the exponential decay model were used to investigate differences in the total increase after the midline and the delay to the maximal increase, through pairwise t-tests. Regression modelling of FJE variability used a quadratic transformation of the sequence length analogous to the study of visual receptive field size (Albright and Desimone, 1987).

### Experiment 1 Results

The effects of motion history on the magnitude of the flash jump effect (FJE) were investigated using a linear mixed effects (LME) regression model. The motion path (continuing vs terminating) and motion sequence length, represented through the distance travelled from the first potential flash position, were included as factors in the LME model. Additionally, a variable intercept for each participant was included to account for individual variation in the magnitude of the FJE. The overall LME was a significant improvement on the null model (p = 1.41e-75), and the specific coefficient results are summarized in table 1 and further discussed below.

**Table 1:** Experiment 1 Regression Model Fitting Results.

| FJE Magnitude Initial Fit |  |  |  |  |
| --- | --- | --- | --- | --- |
| Factor | Fixed Effect ± SE | t-value (df) | p-value | Random Effect STD |
| Intercept | $1.495 \pm 0.163$ | 9.18 (502) | $1.11\text{e-}18$ | 0.746 |
| Terminating Condition | $-1.084 \pm 0.128$ | -5.18 (502) | $3.14\text{e-}16$ | 0.571 |
| Sequence Length | $-0.058 \pm 0.011$ | -8.45 (502) | $3.27e^{-7}$ | N/A |
| FJE Magnitude Midline Split Fit |  |  |  |  |
| Intercept | $1.823 \pm 0.215$ | 8.48 (453) | $3.14e^{-16}$ | 0.750 |
| Terminating Condition | $-1.085 \pm 0.129$ | -8.41 (453) | $5.42e^{-16}$ | 0.564 |
| Sequence Length | $-0.112 \pm 0.024$ | -4.59 (453) | $5.81e^{-6}$ | N/A |
| Midline Resurgence | $0.294 \pm 0.121$ | 2.44 (453) | $1.52e^{-2}$ (0.0152) | N/A |
| FJE Precision Initial Fit |  |  |  |  |
| Intercept | $0.522 \pm 0.034$ | 15.52 (502) | $1.13e^{-44}$ | 0.156 |
| Terminating Condition | $-0.189 \pm 0.02$ | -9.39 (502) | $2.06e^{-19}$ | 0.086 |
| Sequence Length | $-0.0024 \pm 0.0021$ | -1.14 (502) | 0.253 | N/A |
| FJE Precision Quadratic Fit |  |  |  |  |
| Intercept | $0.490 \pm 0.033$ | 14.77 (502) | $2.95e^{-41}$ | 0.156 |
| Terminating Condition | $-0.189 \pm 0.02$ | -9.39 (502) | $2.06e^{-19}$ | 0.086 |
| Fixation-Centered Distance<sup>2</sup> | $-0.00412 \pm 0.00097$ | 4.25 (502) | $2.53e^{-5}$ | N/A |

While the FJE decreases as the flash occurs later in the motion sequence, there was a resurgence in the illusory mislocalization from the flash occurring as the sequence crossed the vertical meridian. We investigated this increase by comparing the minimum FJE before the midline to the maximum after the midline for each participant using a paired t-test. There was a significant increase after the flash had crossed the vertical meridian for both the continuing (t(23) = 4.343, p = 2.396e-4, µ = 0.825 ± 0.393 dva) and terminating conditions (t(23) = 4.317, p = 2.555e-4, µ = 0.508 ± 0.243 dva).

Given the resurgence in the FJE across the midline, we removed the midline from the regression model and included an indicator term for flashes occurring before the midline. The indicator significantly improved the model relative to simply removing the midline (p = 0.0148, AIC change: -3.94). Additionally, the indicator term explaining a significant amount of variation in the FJE (p = 0.0152). In summary, the magnitude of the FJE generally decreases as the length of the motion sequence preceding the flash increases. The reduction in FJE, together with the sudden resurgence across the midline, indicate that motion processing within a hemisphere facilitates more accurate flash localization, while hemifield switching is detrimental.

To investigate if the precision of flash localization also varies with motion history, we fit an LME to the FJE precision with the motion path and motion sequence length as factors as in the previous section; this provided a significant improvement over the null model (p = 9.12e-62). Accounting for individual effects, there was significantly lower variation in the flash-terminating condition (p = 2.06e-19), but no effect of motion sequence length (p = 0.253). We also tested whether positional effects could be due to a symmetric, quadratic relationship across the midline, and found a significant effect of across the squared distance from the midline (p = 2.53e-5).

### Experiment 1 Discussion

In experiment 1, we have demonstrated the FJE observed by Saini and colleagues (2021) is modulated by the motion history before the flashed bar. Specifically, longer preceding motion sequence lengths result in more accurate reports of the position of the flashed bar; this is analogous to the findings of Vreven and Verghese (2005) in the flash-lag effect. The increasing accuracy was present in both the flash-terminating and flash-continuing motion paths. Modulations in positional accuracy could be due to increased visual acuity at lower eccentricities of the flashed bar. However, this would result in symmetric reduction in the FJE centered under the fixation position which was not observed. It is more likely that there is adaptation to the consistent motion of the bar before the colored bar is presented, where longer motion histories result in greater adaptation. While this interpretation conflicts with the conclusions of Coffey and colleagues (2019), results from both studies indicate a stronger motion adaptation result for short motion histories (> 1 s) and are analogous to work in the flash-lag effect (Cantor and Schor, 2007). The transient increase in FJE near the vertical meridian may be due differences in adaptation between hemispheres, or a switch from foveofugal to foveopetal motion which has been shown to modulate the flash-lag effect (Kanai et al., 2004).

### Experiment 2 Results

The goal of experiment 2 is to test the hypothesis that the findings from experiment 1 are the result of hemisphere-dependant motion habituation that is reset for motion crossing the vertical midline. We explored a wider range of motion sequence lengths before the vertical midline by varying the fixation position to the left, right, and center of the screen, thus creating longer and shorter motion sequences before crossing the midline. If the decrease in FJE for greater motion sequence lengths is the result of habituation of the motion of the bar within a given hemisphere, then the decrease should continue for flashes occurring within a hemifield, regardless of the position of the fixation. Additionally, if the increase in FJE across the midline is the result of habituation in one hemifield not transferring to the other hemifield, then it should be independent of the fixation position. However, if the increase is a result of an allocentric cue for the center of the display, the increase should occur at the same absolute position regardless of the change of fixation position and the visual vertical meridian. Compared to experiment 1, experiment 2 added a potential flash position to the beginning and end of the sequence and compared a central fixation with a fixation 3.5 dva earlier or later in the motion sequence.

Given the resurgence of the FJE upon crossing the midline in experiment 1, we first checked for this resurgence for the motion terminating condition for each of the three fixation positions. There was a significant increase in the FJE magnitude across the midline as in experiment 1 for each fixation position and motion termination condition (Table 2). Following from experiment 1, we fit an LME model to the FJE magnitude using the sequence length, motion termination condition, midline indicator (removing points on the midline), and an interaction between the sequence length and fixation position as factors (Table 3). This model performed significantly better than a null model (p = 1.47e-201). While the length of the sequence alone did not influence the FJE magnitude, there was a significant interaction with the fixation position. We speculated that this may be due to an adaptation effect, like that observed in experiment 1, centered on the horizontal meridian that moves with the fixation positions.

**Table 2.** Experiment 2 Midline Resurgence Comparison. Results of a t-test of signed rank test (indicated by $), reported as the mean change (p-value) from the minimum value before to maximum after the horizontal meridian.

|  | Flash-Terminating | Flash-Continuing |
| --- | --- | --- |
| Early Fixation | +1.37 dva ($2.29\text{e}^{-5}$) \$ | +2.81 dva ($2.89\text{e}^{-5}$) |
| Middle Fixation | +1.39 dva ($1.23\text{e}^{-6}$) | +2.63 dva ($1.30\text{e}^{-7}$) |
| Late Fixation | +1.22 dva ($4.47\text{e}^{-7}$) | +0.71 dva ($5.39\text{e}^{-3}$) |

**Table 3:** Experiment 2 Regression Model Fitting Results.

| FJE Magnitude Initial Fit |  |  |  |  |
| --- | --- | --- | --- | --- |
| Factor | Fixed Effect ± SE | t-value (df) | p-value | Random Effect STD |
| Intercept | $1.537 \pm 0.441$ | 3.48 (1603) | $5.056\text{e}^{-4}$ | 1.105 |
| Sequence Length | $-0.014 \pm 0.032$ | -0.44 (1603) | 0.6607 | N/A |
| Fixation Position | $0.217 \pm 0.032$ | 6.73 (1603) | $2.412\text{e}^{-11}$ | 1.044 |
| Terminating Condition | $-2.024 \pm 0.224$ | 9.04 (1603) | $4.296\text{e}^{-19}$ | N/A |
| Midline Resurgence | $0.462 \pm 0.138$ | 3.34 (1603) | $8.721\text{e}^{-4}$ | N/A |
| Sequence Length*Fixation Position | $-0.011 \pm 0.003$ | -4.09 (1603) | $4.577\text{e}^{-5}$ | N/A |
| FJE Flash-Terminating |  |  |  |  |
| Intercept | $0.0004 \pm 0.376$ | 0.001 (788) | 0.999 | 0.429 |
| Sequence Length | $-0.064 \pm 0.031$ | -2.06 (788) | 0.039 | N/A |
| Fixation Position | $0.039 \pm 0.031$ | 1.26 (788) | 0.208 | N/A |
| Midline Resurgence | $-0.198 \pm 0.134$ | -1.48 (788) | 0.139 | N/A |
| Sequence Length*Fixation Position | $0.0034 \pm 0.0025$ | 1.37 (788) | 0.170 | N/A |
| FJE Precision Quadratic Fit |  |  |  |  |
| Intercept | $2.019 \pm 0.177$ | 11.39 (1845) | $4.222\text{e}^{-29}$ | 0.837 |
| Time-to-Fixation<sup>2</sup> | $0.0063 \pm 0.0011$ | 5.61 (1845) | $2.328\text{e}^{-8}$ | N/A |
| Terminating Condition | $-0.982 \pm 0.224$ | -8.27 (1845) | $2.459\text{e}^{-16}$ | 0.483 |
| Time-to-Fixation<sup>2</sup>* Terminating Condition | $0.0027 \pm 0.0016$ | 1.71 (1845) | 0.0876 | N/A |

We separately characterized the adaptation effect for the flash-continuing and terminating conditions because of the differences in the pattern of adaptation. An LME was fit to only the flash-terminating condition (Figure 4B) and found a significant decrease in FJE across the length of the sequence (p = 0.0394) that was unchanged for the different fixation positions. However, a general resurgence was not significant across the fixation positions (p = 0.139).

**Figure 1.**
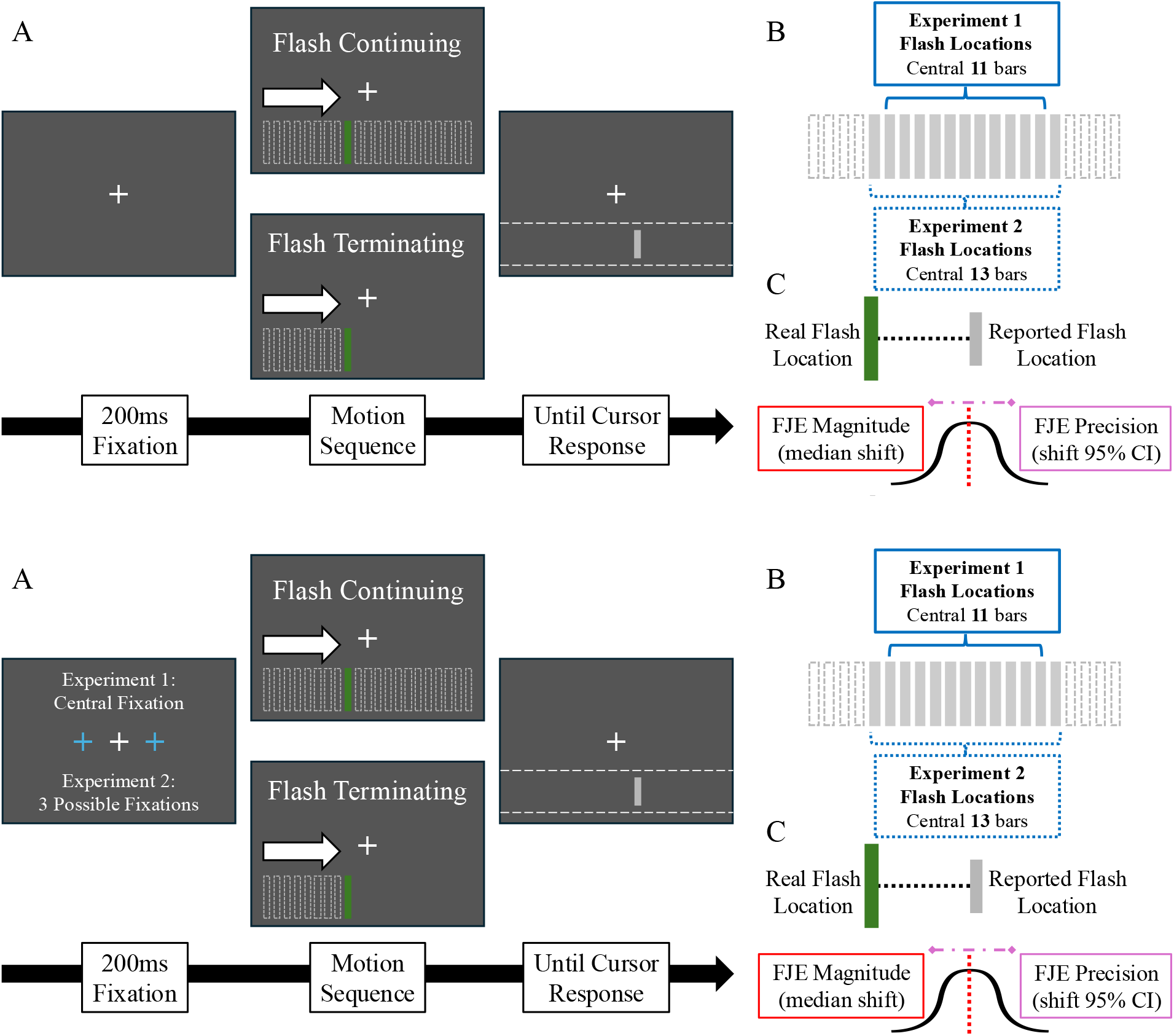
Experimental Methods. A) Progression of an experimental trial began with a 200ms fixation on a cross at the center of the display (Exp 1), or center, 4 dva to the left, or right of center (Exp 2). Following the initial fixation period, grey bars began an apparent motion sequence from the left or right of the display (only rightwards motion depicted). One of the bars in the motion sequence was instead presented as a color (blue, red, yellow, or green in Exp 1, only green in Exp 2). The colored bar was either presented as the last in the sequence (flash-terminating) or with subsequent grey bars extending to the other edge of the display (flash-continuing). After the sequence ended, participants then indicated where they observed the colored flash with a vertically locked probe bar. B) The colored flash could be presented at any of the central 11 bars (Exp 1) or 13 bars (Exp 2), with the overall length of the sequence unchanged in the flash-continuing condition. C) The magnitude of the flash-jump effect (FJE) was computed as the offset between the reported flash position and the actual flash position, corrected for the direction of motion (i.e. further in the direction of motion was positive while earlier was negative). The median FJE for each participant and experimental condition was used to assess FJE magnitude, while a 95% confidence interval for each participant and condition was used to assess the precision of the FJE.

**Figure 2.**
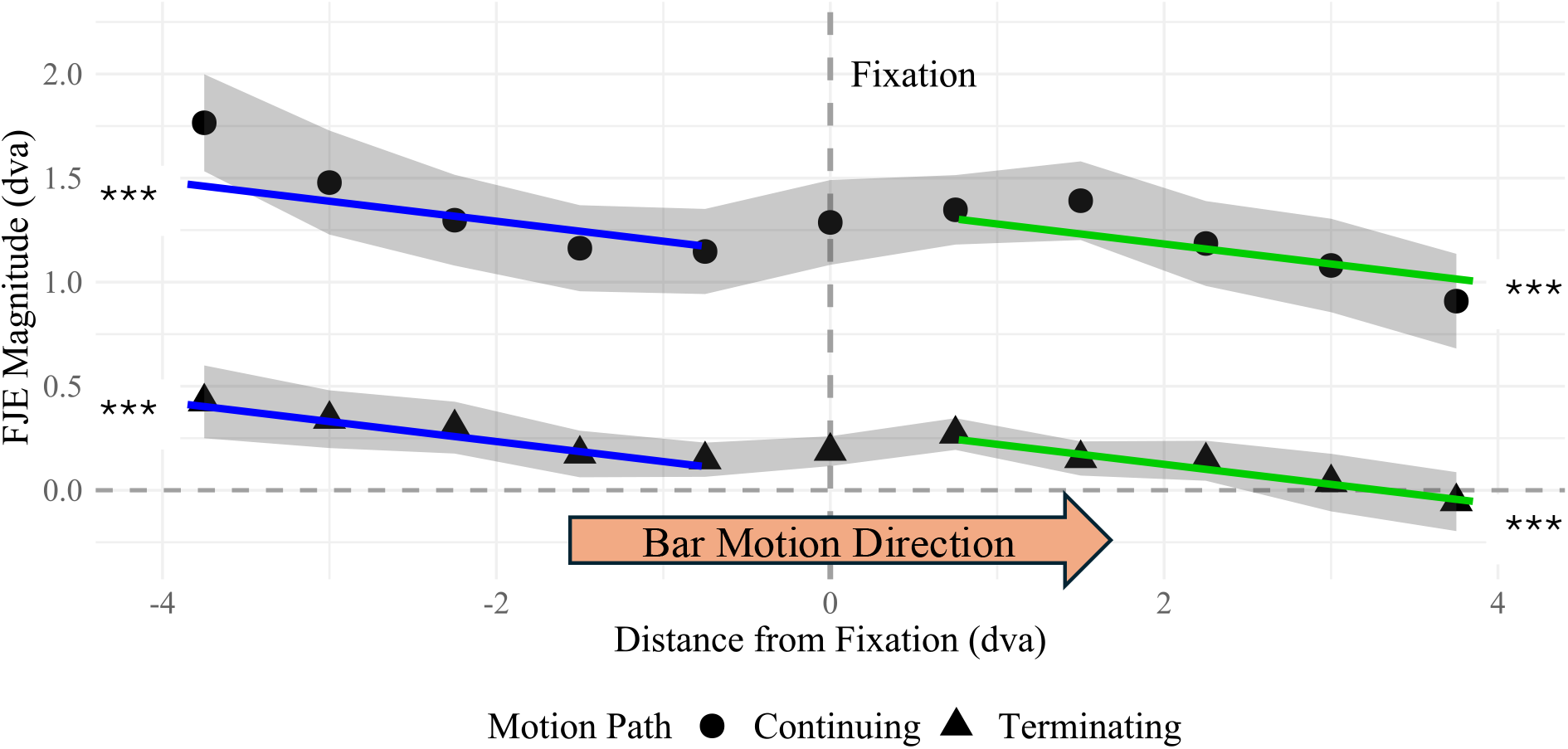
Experiment 1 Flash Jump Effect Magnitude Over Flash Positions. FJE magnitude plotted across the horizontal distance of the colored bar flash from the fixation cross, collapsed across motion direction and depicted as rightward motion. Each point represents the mean +/- SEM shift for each participant median in the flash-terminating (triangles) and flash-continuing (circles) motion conditions. Greater positive values indicate less accurate reports of the flash position further along its motion path. The lines are the fit values of different conditions in a mixed effects regression model across the length of the motion sequence. The blue lines are the fits for FJE magnitude before crossing the horizontal meridian, while the green lines are the fits for flashes after the horizontal meridian. ***: p < 0.001 for the different sections of the mixed effects model.

**Figure 3.**
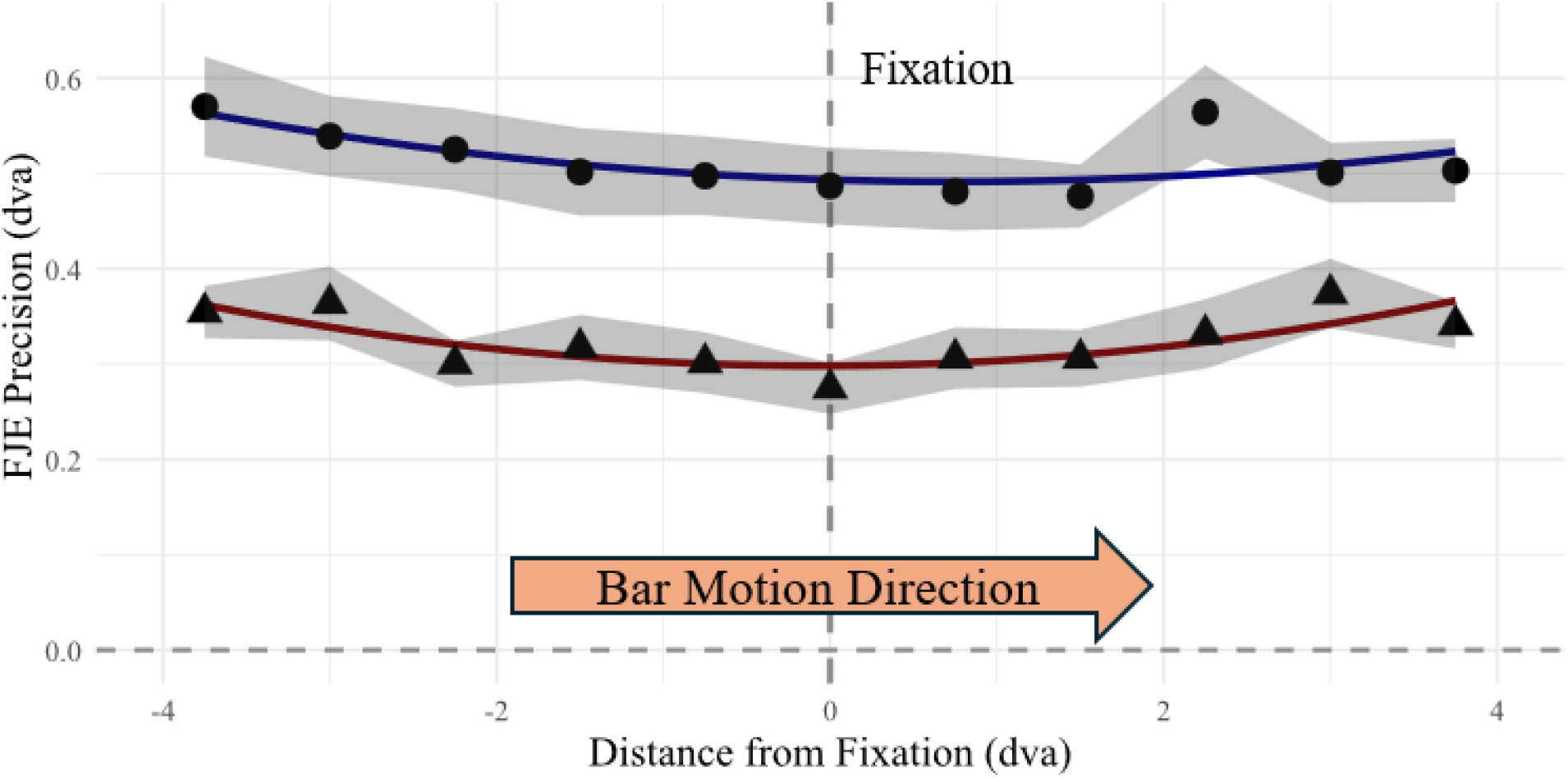
Experiment 1 Flash Jump Localization Precision Over Flash Positions. FJE precision plotted across the horizontal distance of the colored bar flash from the fixation cross, collapsed across motion direction and depicted as rightward motion. Each point represents the mean +/- SEM 95% confidence interval range for each participant in the flash-terminating (triangles) and flash-continuing (circles) motion conditions. Higher values indicate greater variability in the reporting of flash position.

**Figure 4.**
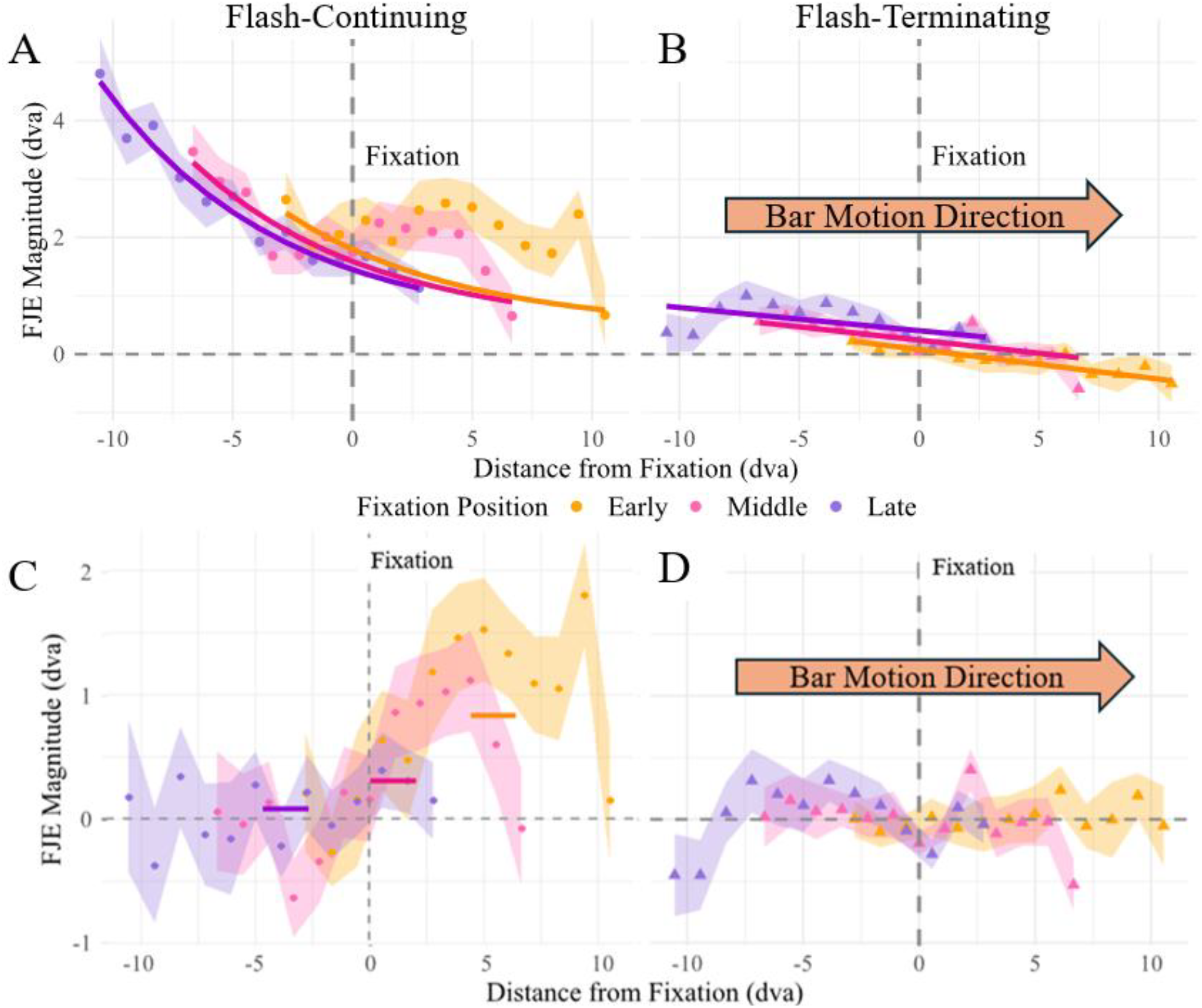
Experiment 2 Flash Jump Localization Accuracy Over Flash and Fixation Positions and Fit Residuals. The magnitude of the FJE in the flash continuing (A/C) and terminating (B/D) motion path conditions across the 3 fixation position times: early (yellow), middle (pink) and late in the motion sequence (purple). A/B) The points and shaded regions represent the mean ± SEM for the motion path conditions. The solid lines represent the best regression fits to the data. Note that the exponential decay in A was fit only to the data before crossing the fixation (details in text). C/D) The residual FJE magnitude after correcting for the predicted values of the fits in A/B. The solid bars represent the mean residual value and are plotted at the distance from fixation that aligns with the average maximum residual value, reflecting the resurgence in FJE after crossing the midline.

**Figure 5.**
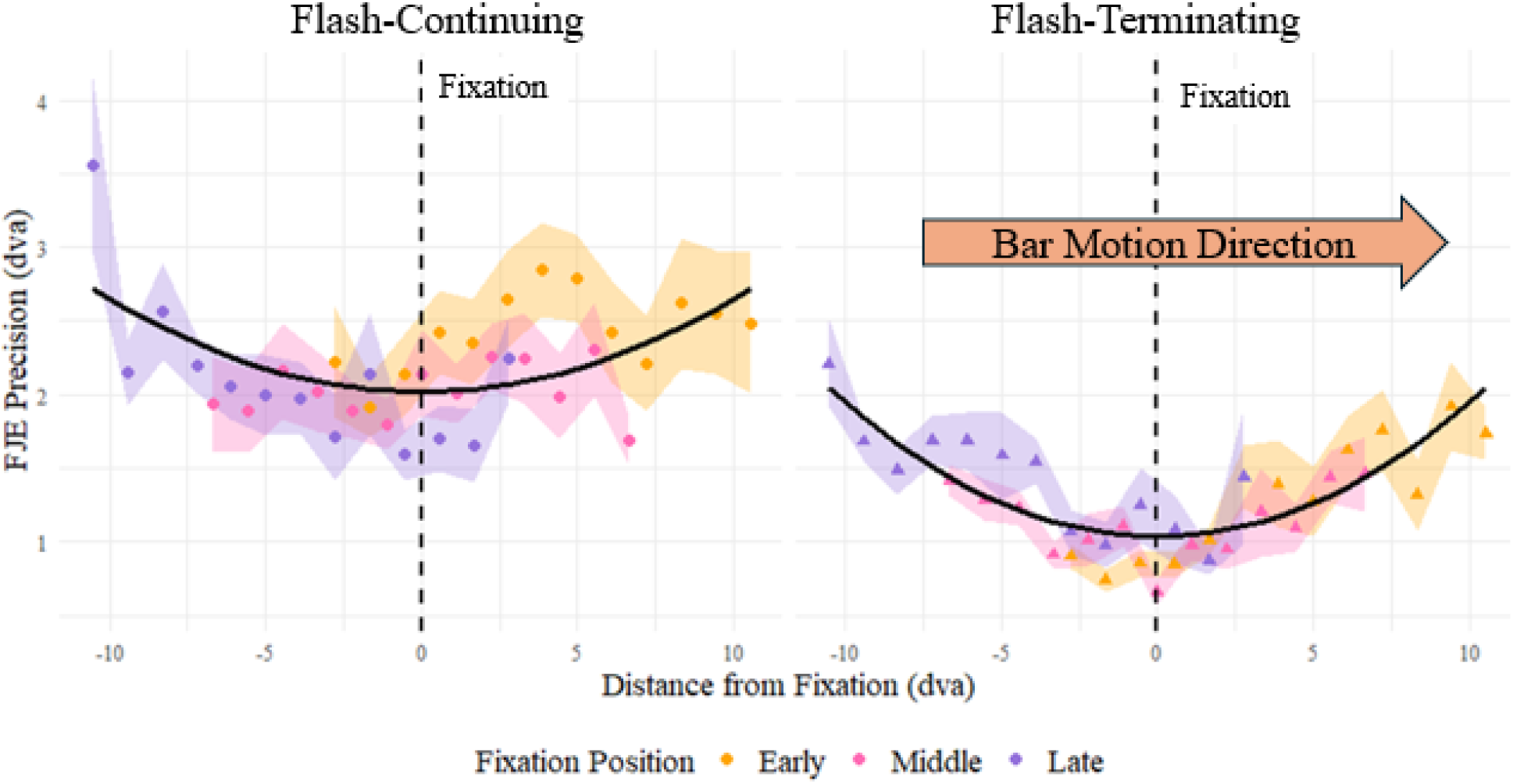
Experiment 2 Flash Jump Localization Precision Over Flash and Fixation Positions. The FJE precision plotted across the horizontal distance of the colored bar flash from the fixation cross, following the same conventions as Figure 3. Note that the FJE precision has been collapsed across motion direction and depicted as rightward motion and higher values indicate greater variability in the reporting of flash position. Each point represents the mean +/- SEM 95% confidence interval range for each participant in the flash-terminating (triangles) and flash-continuing (circles) motion conditions across the 3 fixation positions presented early (yellow), middle (pink) or late (purple) in the motion sequence.

For the flash-continuing condition, we compared a linear fit with a more biologically plausible exponential decay model of motion adaptation (Priebe et al., 2002). This was initially done using only flashes before the midline and the latest occurring fixation position to allow for the most extensive characterization of the motion adaptation signal without hemifield interference (Figure 4A). The exponential decay model performed much better than the linear model (AIC change: - 3.014), and resulted in the following model:

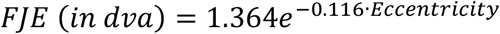

With a participant-wise intercept of 0.011 ± 0.55 dva, indicating substantial variation in the overall participant FJE but a general lower bound for adaptation that is close to the accurate perception of the flashed bar. Using this fit, the same decay function and parameters were used to fit an adaptation model to the middle and shortest paths before the midline alongside an additional parameter allowing translational shifts along the sequence (i.e. the d parameter in: *a*·e^(-*b*·(Eccentricity+d))^ + *c*). These fits indicated that the exponential decay was a good fit for the FJE magnitude at all 3 fixation positions relative to a linear model (Middle AIC change: -7.962; Short AIC change: -1.385), without significant evidence in a shift of the decay function across the 3 fixation points (all p > 0.064).

Having characterized the adaptation occurring before the midline, we investigated the resurgence after its crossing through the FJE after removing the FJE predicted by the exponential decay model. From the residual FJEs, we identified the maximum value and when it occurred after the midline for each participant (Table 4 | Figure 4C/D). We found that the resurgence was significantly above the predicted decay value for all fixations (all p < 6.88e-5), but with increasing resurgence strengths the earlier the midline was crossed. Additionally, the timing of the peak resurgence varied by the length of the motion path before crossing the midline in a pattern that aligned with the difference in timing of the fixations. That is, the resurgence was stronger the earlier in the motion sequence that the bar crossed the midline. However, the timing of the peak occurred at a similar time in the overall path of the bar, independent from the fixation position. Together these results show that prior to the vertical meridian, the magnitude of the flash-jump illusion follows an exponential decay; then upon crossing the meridian, the illusion has a resurgence which increases based on the length of the motion sequence prior to the vertical meridian of the display. Thus, the more habituation that occurs before crossing the midline leads to more recovery of the flash jump illusion after crossing it.

**Table 4:**
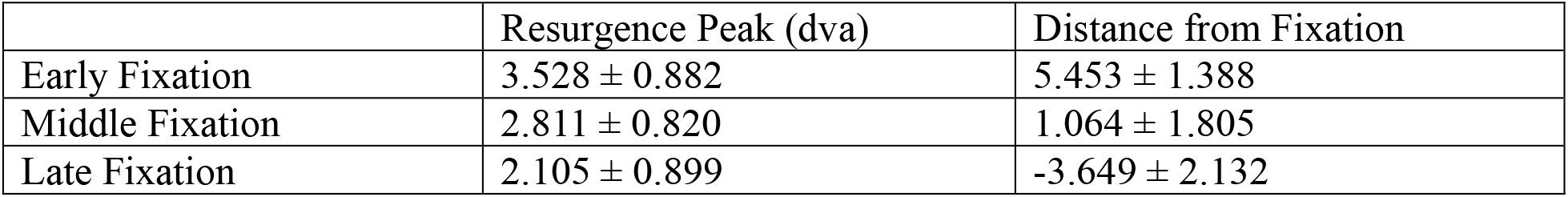
Experiment 2 resurgence peaks after correcting for motion adaptation for fixation positions and the time that the peak occurred after the fixation. Presented as mean ± 95% CI.

The precision of the FJE was determined through 95% confidence intervals for each participant and condition and was investigated using the same analysis as experiment 1: a quadratic LME across flash positions centered on the fixation position (Table 3). This model performed significantly better than a linear fit on the same data (AIC change: -92.61). The prediction of the location of the bar was significantly more precise in the flash-terminating condition, replicating the results of experiment 1. Precision increased for bars closer to the midline and was not significantly different between motion terminating conditions (p = 0.0876). This demonstrates that the relationship between flash location and FJE precision is independent of the terminating condition and is generally more precise for bars near fixation.

### Experiment 2 Discussion

In experiment 2, we replicated the adaptation effect for increased motion sequence length and the transient increase in FJE as the bar crosses hemifields observed in experiment 1. By moving the position of the fixation cross, but maintaining a constant head position, we were able to test whether the motion adaptation occurs locally within a hemisphere or globally across the visual field through the length of the motion sequence at the time of the flash. The FJE, and other motion-induced position shifts, have been suggested to result from the integration of an accurate perception of the position of the flash with an extrapolated position based on the motion of the bar (Sundberg et al., 2006; Khoei et al., 2017; van Heusden et al., 2018; Saini et al., 2021; Keil et al., 2022). We have demonstrated that adaptation of these motion signals is restricted within a hemisphere, indicating that the signals underlying the FJE likely originate in early visual areas without large bilateral receptive fields. The FJE has been recorded in area V4 (Sundberg et al., 2006), where receptive fields have limited ipsilateral representation (Desimone et al., 1993; Tootell et al., 1998). Similarly, the motion sensitive area MT has only limited ipsilateral representation (Tootell et al., 1998; Huk et al., 2002; Amano et al., 2009), and its activity has been heavily implicated in the presence and strength of the similar flash-lag illusion (Maus et al., 2013a/b; Wang et al., 2022).

## General Discussion

Across 2 experiments and experimental settings, we used the flash-jump illusion: a mislocalization of a flashed bar further along its motion path that depends on motion continuing after the flash (Sundberg et al., 2006; Saini et al., 2021) to show that longer preceding sequences before the flash resulted in reduced forward mislocalization, and more accurate perceptions of the flash location. We attribute this effect to adaptation to the consistent motion of the bar leading to decreased motion extrapolation. Notably, the motion adaptation effect occurred regardless of whether motion after the flash was present but was more prominent when the motion sequence continued. We hypothesized that the adaptation effect occurred locally within a hemisphere due to the transient increase in flash-mislocalization when crossing the horizontal meridian. By shifting the position of the visual midline, we confirmed that the adaptation effect is localized within a hemifield and transfers only minimally to the opposite hemisphere, consistent with evidence that regions near the horizontal meridian are represented in both hemispheres (Fendrich et al., 1996). Notably, the level of adaptation maintained across the midline increased for longer motion sequences preceding the hemifield switch; that is, a smaller FJE increase was observed immediately after the midline for longer sequences.

The flash-jump illusion studied here is an extension of the original flash-lag paradigm (Nijhawan, 1994) that employed 2 separate stimuli: a rotating bar and a stationary bar that appeared briefly adjacent to the moving bar. Subjects report the static stimulus as occurring behind the moving stimulus, perceiving it as lagging the moving stimulus. Instead of a spatial judgement of physically separate stimuli, the flash-jump illusion instead addresses the integration of a spatial judgement with a non-spatial feature of the stimulus, often color (Cai and Schlag, 2001; Sundberg et al., 2006; Saini et al., 2021). Theories explaining motion-induced position shifts have evolved over time (Hogendoorn 2020), with the motion extrapolation theory retaining substantial support. Motion extrapolation supposes that the brain develops an internal representation of an object’s motion over time to facilitate better tracking and action towards moving targets (Maus and Nijhawan, 2008; for a review see Hogendoorn, 2020). In the context of the motion induced position shifts, the extrapolation of a moving object causes it to be perceived further along its trajectory and ahead of adjacent stationary stimuli.

### Alternatives to Motion Extrapolation

There are, however, alternative theories of motion perception that do not require an internal model of motion to be formed, and we briefly address prominent such theories in this section. The flash-jump paradigm makes use of a featural change in a single object, therefore theories like the attentional shifting hypothesis (Baldo and Klein, 1995) that induce attentional mechanisms relating to the presence of a 2^nd^ object in the flash-lag effect cannot explain the flash-jump effect. The differential latencies hypothesis (Whitney and Murakami, 1998) proposes that the moving bar is processed faster than static objects or events like the colour change of the bar. To incorporate the adaptation effect observed here, the processing speed of the flash would have to increase with motion history, or slow the processing speed of the moving bar, or some combination of both amounting to a >200ms overall change over the motion history. Similarly, the discrete sampling hypothesis (Schneider, 2018) posits that the path of the bar is collapsed into time windows where position estimates are taken from the latest point in each window. The width of the windows would need to decrease with longer motion history, allowing for more accurate flash localization; however, neither the discrete sampling or differential latency hypotheses readily explain a sudden resurgence in FJE across the midline. The postdiction or temporal integration theory (Brenner and Smeets, 2000; Eagleman and Sejnowski, 2000, 2007) supposes that information is collected after the flash and biases the localization further along the motion sequence. However, the range of this postdiction window has been estimated at 60-80ms (Brenner and Smeets, 2000; Eagleman and Sejnowski, 2007), which is much shorter than the overall perceived shift, and still does not readily allow for shifts in the range of the window based on the preceding motion sequence. Thus, none of these alternate theories can easily or readily explain how motion history affects the magnitude of the mislocalization.

### Adaptation in the Context of Motion Extrapolation

We suggest that the adaptation effect observed in our study is due to neural habituation affecting the magnitude of the motion extrapolation signal.in motion selective area MT. This is supported by the magnitude of the flash-lag effect being increased or decreased through TMS (Maus et al., 2013) and tDCS (Wang et al., 2022) over MT+. Given that the adaptation we observed was limited within a hemifield, it is unlikely that MST is the source of the adaptation due to that area’s large bilateral receptive fields (Raiguel et al., 1997; Huk et al., 2002). In contrast, the receptive fields in MT show only limited representation of the ipsilateral visual field (Raiguel et al., 1995), consistent with our results. Further, adaptation of MT has been shown to reduce perceived speed (Krekelberg et al., 2006), which suggests that speed processing being reduced through adaptation may be the driver for the reduction in motion extrapolation.

The transient increase in FJE at the visual midline can be explained through the integration of motion extrapolation signals from the habituated and unhabituated hemispheres. That is, once the bar moves into a region of the visual field that is represented bilaterally, 2 extrapolation signals become available: from the area MT that processed the motion path before bilateral representation and has habituated, and from the other area MT where the bar’s motion is just appearing with little to no habituation. Motion extrapolation has been modelled previously through Bayesian inference where the motion history of an object is used as a prior to determine the actual position of the flashed object. The Bayesian model of motion extrapolation has been used previously to describe the effect of the flash-jump illusion (Sundberg et al., 2006; Saini et al., 2021). Saini and colleagues (2021) suggested that the color modulation of FJE magnitude in their study was due to differential weighting of the V4 prior observed by Sundberg and colleagues (2006). We propose that both left and right area MT extrapolation signals are needed as individual priors to determine the position of the flashed bar around the midline where a bilateral representation is available. The weight of each prior is derived from the firing rate of the MT neurons, such that greater adaptation leads to lower firing rates and a higher bayesian weight. From our results, the shortest motion sequence length before the midline led to the greatest resurgence in FJE reflecting a lower weight of the pre-midline motion extrapolation prior. This interpretation supports the basic Bayesian principle that predictions are based on the most reliable information.

The question remains: where in the visual hierarchy are the color-change and motion extrapolation signals integrated? The primary motion extrapolation signals originate in MT, while the representation of the colored flash is processed in V4, both of which have strong hemifield separation. Given the known connections between V4 and MT (Maunsell and van Essen, 1983; Ungerleider et al., 2008), the most parsimonious site for the initial binding of the flash and moving bar is MT itself. However, this integration alone could not adequately explain our finding of the resurgence in the adaptation effect. Instead of observing a consistent magnitude and timing for the peak resurgence, as would be expected from a clear hemispheric separation, we observed a smoother transition between one adaptation signal to another. As in the last section, the combination of 2 adapting motion extrapolation signals, combined downstream of the initial binding in MT, provides a better explanation.

We thus propose that MST is the site of our observed resurgence, acting as an integrator of the adapting signals from the upstream MTs in both hemispheres. MST receives inputs from MT and consists of neurons with large, bilateral receptive fields (Raiguel et al., 1997; Amano et al., 2009). In our explanation, MST acts as an integrator of motion signals from MT, weighting them based on their reliability to create a persistent representation of the motion of the bar. The adaptation in MT thus serves as an indicator of reliability.

### Hemispheric Processing and Object Tracking

Our findings also have implications for perceptual estimates of speed and object tracking. The accuracy of tracking an object that crosses between hemifields is decreased relative to those that stay within a hemifield (Alvarez and Cavanaugh, 2005), a cost attributed to early sensory processing (Minami et al., 2019). Our results indicate that the within hemifield tracking advantage is likely due in part to separate adaptation to the moving objects in each hemisphere. This perceptual limitation may serve as a diver of an important oculomotor behaviour: smooth pursuit eye movements. By pursuing a moving object, an observer stabilizes its position on the fovea, ensuring that it is represented continuously by both hemispheres (Nijhawan, 2001). Smooth pursuit eye movements would then prevent the abrupt crossover between the adapted and unadapted hemisphere, leading to a more stable perception, free from the resurgence in positional errors that we observed.

This distinction may reflect the different goals of ‘vision for perception’ and ‘vision for action’ (Goodale and Milner, 1992). Our fixation-based task probes only the perceptual system, highlighting the circuitry via the limitations like the perceptual cost of crossing the midline. This framework is supported by recent findings demonstrating that different motor actions respond different to motion illusions. Specifically, work from Lisi and Cavanagh (2024) found that interceptive saccades are less influenced by illusory motion than smooth pursuit or fixated perceptual reports. However, they also found that smooth pursuit eye movements more closely resembled the biases found in fixated perceptual reports, suggesting that perception and pursuit share common motion representations (Lisi and Cavanagh, 2024). This dissociation offers a more nuanced view of the perception-action model. Rather than a strict dichotomy, visual processing may operate on a continuum of behavioural goals, supported by the hierarchy of neural timescales (Murray et al., 2014; Lisi and Cavanagh, 2024). On one end, rapid interceptive saccades represent purely action-based perception, relying on motion signals extrapolated only over short intervals (Lisi and Cavanagh, 2024). On the other end, fixated perception may integrate longer timescales to provide more stable, detailed representations. Note that this distinction in integration or extrapolation times is across the goals of perception, rather than an account for the effects of motion extrapolation itself covered earlier. Smooth pursuit falls somewhere between, making use of longer integration windows like fixated perception (Lisi and Cavanagh, 2024), yet serves the action-oriented goal of stabilizing a moving stimulus. We propose that the MST is well positioned to integrate motion signals flexibly, selectively weighting inputs based on the temporal integration required for the current goal. This notion is supported by the representation of self motion in MST, but not MT neurons (Inaba et al., 2007), allowing MST to inform smooth pursuit even when it would require head or body motion.

## Conclusion

By examining the flash-jump effect across various motion paths, we have demonstrated that motion adaptation and motion extrapolation occur independently in each hemifield, influencing the perception of an object’s position. We argue that this phenomenon is best explained by a motion extrapolation mechanism that habituates within the hemispherically separated areas of MT. The resurgence of the illusion at the visual midline reflects a Bayesian integration of signals from both the adapted and unadapted hemispheres, a process likely occurring in area MST. These findings, observed during fixation, reveal a fundamental constraint of the ‘vision for perception’ system with moving stimuli. More importantly, they highlight the functional significance of the ‘vision for action’ system, suggesting that the drive to smoothly track an object with our eyes is a behavioural solution to overcome perceptual discontinuities near the midline. Our work solidifies motion extrapolation as the key mechanism underlying these motion-induced illusions and reframes the cost of interhemispheric transfer as a driver of oculomotor behavioural strategies.

